# Polymer Stiffness Regulates Multivalent Binding and Liquid-Liquid Phase Separation

**DOI:** 10.1101/2020.05.13.093245

**Authors:** E. Zumbro, A. Alexander-Katz

## Abstract

Multivalent binding is essential to many biological processes because it builds high affinity bonds by using several weak binding interactions simultaneously. Multivalent polymers have shown promise as inhibitors of toxins and other pathogens, and they are important components in the formation of biocondensates. Explaining how structural features of these polymers change their binding and subsequent control of phase separation is critical to designing better pathogen inhibitors and also to understanding diseases associated with membraneless organelles. In this work, we will examine the binding of a multivalent polymer to a small target. This scenario could represent a polymeric inhibitor binding to a toxic protein or RNA binding to an RNA-binding protein in the case of liquid-liquid phase separation. We use simulation and theory to show that flexible random-coil polymers bind more strongly than stiff rod-like polymers and that flexible polymers nucleate condensed phases at lower energies than their rigid analogues. We hope these results will provide insight into the rational design of polymeric inhibitors and improve understanding of membraneless organelles.

**Statement of Significance:** Multivalent polymers are essential for many biological systems, including targeting pathogens and controlling the formation of liquid-liquid phase separated biocondensates. Here, we explain how increasing polymer stiffness can reduce multivalent binding affinity to a small target such as a toxic protein and how modulating polymer stiffness can change the phase boundary for liquid-liquid phase separation. These results have implications for designing stronger pathogen inhibitors and provide insights on neurodegenerative diseases associated with abnormal biocondensate formation.

## Introduction

Multivalent binding interactions are commonly found throughout biology and synthetic applications. These interactions use multiple weak binding sites to simultaneously bind to another species. Using many low-affinity binding events simultaneously enhances the overall binding affinity much more than the sum of the constituent monovalent binding interactions [1]. Multivalent binding can take on many different geometries and previous research has been done on nanoparticles, sheets, dendrites, and polymers for numerous applications [2, 3]. In this work, we will focus on multivalent polymers as they pertain to toxin inhibition along with implications for nucleating liquid-liquid phase separation in biocondensates.

Synthetic multivalent polymers have shown promise at binding to and inhibiting multivalent sugar-binding proteins called lectins [4–9]. Monovalent sugar-protein binding affinities are typically weak, in the millimolar to micromolar range, so multivalency is essential to creating high binding affinities [9, 10]. Binding to lectins is an exciting avenue for combatting infection because many toxins are lectins such as cholera toxin, shiga toxin, and others that cause diarrheal diseases, and because bacteria and viruses use lectins on their surface to bind to the glycocalyx on our cells [2, 11]. Mucins, the megadalton weight glyprotein polymer found in mucus, are thought to use their glycan brushes as binding decoys, exploiting multivalency to bind to pathogenic lectins and prevent infection [12]. Attempts to mimic this capability with synthetic polymers have been successful, but the effects of polymer backbone flexibility and characteristic ratio *C_∞_* on the binding of polymers much larger than their targets has not received theoretical study. Previous studies on the flexibility of multivalent binding have focused on species of similar size binding to each other. In these cases, small molecules or oligomers with binding sites precisely spaced to the target found rigid linkers to minimize the entropic cost of binding, resulting in the highest binding affinity [2,5,8,13,14]. When the polymer chain’s end-to-end distance is on the same scale as the binding target, stiff linkers between binding sites on an antibody and rigid sections near the ligands of a divalent binder were shown to be higher affinity than their flexible counterparts [15,16]. In contrast, very few studies have considered polymers much larger than the size of their targets, even though this is the scale of native mucins and many of the previously mentioned synthetic inhibitors tested experimentally [17, 18]. We anticipate that because, unlike small divalent oligomers, many-valent polymers allow for many binding site pairs with different spacings, large multivalent polymers may benefit from higher flexibility which allows them to sample more binding combinations [1, 3, 19]. Theoretical research on this relevant size scale has not considered the effect of stiffness of the polymer chain and how this controls binding affinity to a small multivalent target. Here, we examine how a single target binds to large many-valent polymers of increasing stiffness and provide a theoretical explanation for the difference in binding modes between a flexible random coil polymer and a stiff wormlike polymer chain.

Understanding multivalent polymers and their binding is also essential to controlling liquid-liquid phase separation in membraneless organelles [20–22]. Research has shown that polymeric binding characteristics such as valency and individual binding site strength can be used to control the phase separation boundary [19, 20, 23]. Other studies have shown that polymer properties indirectly related to binding sites such as solvation volume can determine the difference between a cross-linked gel and a phase separated system [23]. Because of this, we expect that flexibility of the polymer could also be an important factor in controlling liquid-liquid phase separation. Dysregulation of the phase separation in membraneless organelles is a common feature of neurodegenerative diseases like Alzheimer’s, Parkinson’s, and ALS [20, 21, 24], and so investigating how features of multivalent polymers can change the phase boundary are essential. We hope that this research will contribute understanding to how the stiffness of polymer chains can modulate nucleation of condensed phases and thus how changes in polymer stiffness could lead to aberrant condensates or disease.

In this work, we focus on how the change in polymer stiffness modifies its binding affinity to a much smaller target. This scenario could represent a coarse gain model of a polymer binding to lectin in the case of toxin inhibition or a long section of RNA binding to a smaller RNA-binding protein in the case of biocondensates [25,26]. First, we will discuss the case of a single target binding to the polymer and provide a theoretical understanding of how polymer stiffness changes binding affinity. In the second half of the paper, we will present results for many targets simultaneously binding to the polymers for an array of target solubility limits. We explore how polymers can nucleate condensed phases and how polymer stiffness changes this phase boundary. We hope that these results can aid in the study of polymeric inhibitors as well as in the understanding of liquid-liquid phase separation of biocondensates.

## Computational Methods

To study the multivalent binding of polymers with varying stiffness, we used a coarse-grained Brownian dynamics simulation with a bead-spring polymer and a spherical target represented as a single bead of the same size as the polymer beads [18]. We applied Brownian dynamics to each bead governed by the equation:

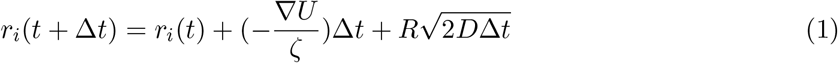

Where *r_i_* is the position of the bead at time *t* in the direction *i* = x, y, or z, *R* is a random number drawn from a normal distribution with a mean of 0 and a standard deviation of 1, *ζ* is the drag coefficient, and *D* = *k*_B_*T*/*ζ* is the diffusion coefficient. The forces each bead experiences due to interactions with the surrounding polymer or target are captured in ∇*U* where *U* is a potential energy that combines contributions from connectivity, bending, excluded volume, and binding. These are added together as *U* = *U*_sp_ + *U*_Bend_ + *U*_LJ_ + *U*_Bind_.

Connectivity along the polymer chain is controlled by harmonic springs with the equation:

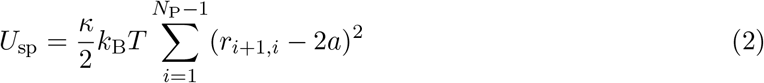

where *r_ij_* is the distance between polymer beads, *a* is the radius of a simulation bead, and *κ* was chosen to be 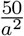, a value sufficiently large enough to prevent the polymer from stretching apart under normal Brownian forces.

To control the flexibility of the polymer chain we use the Kratky-Porod wormlike chain model [28, 29], and introduce an additional spring placed between every next nearest neighbor along the chain. This is a commonly employed scheme in some force fields such as MARTINI [27]. This spring imposes an energetic penalty for bending around the straight/stiff configuration between two bonds as shown in Figure 1. This added spring is implemented using the potential below:

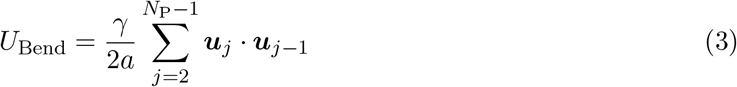

where *γ* is the bending rigidity and 2*a* is the equilibrium length between two bead centers. ***u***_*j*_ is the unit vector pointing from bead *j* to bead *j* + 1 [30, 31]. This allows us to control the persistence length or *C_∞_* by modulating *γ*. When *γ* = 0 we reproduce a freely jointed chain [32], and as *γ* increases, the chain becomes a semiflexible polymer and a stiff, rod-like polymer at high *γ*. The end-to-end distance of the polymer with increasing values of *γ* is shown in Figure 2. The corresponding values of persistence length *p* and characteristic ratio *C_∞_* are also displayed for reference.

**Figure 1:**
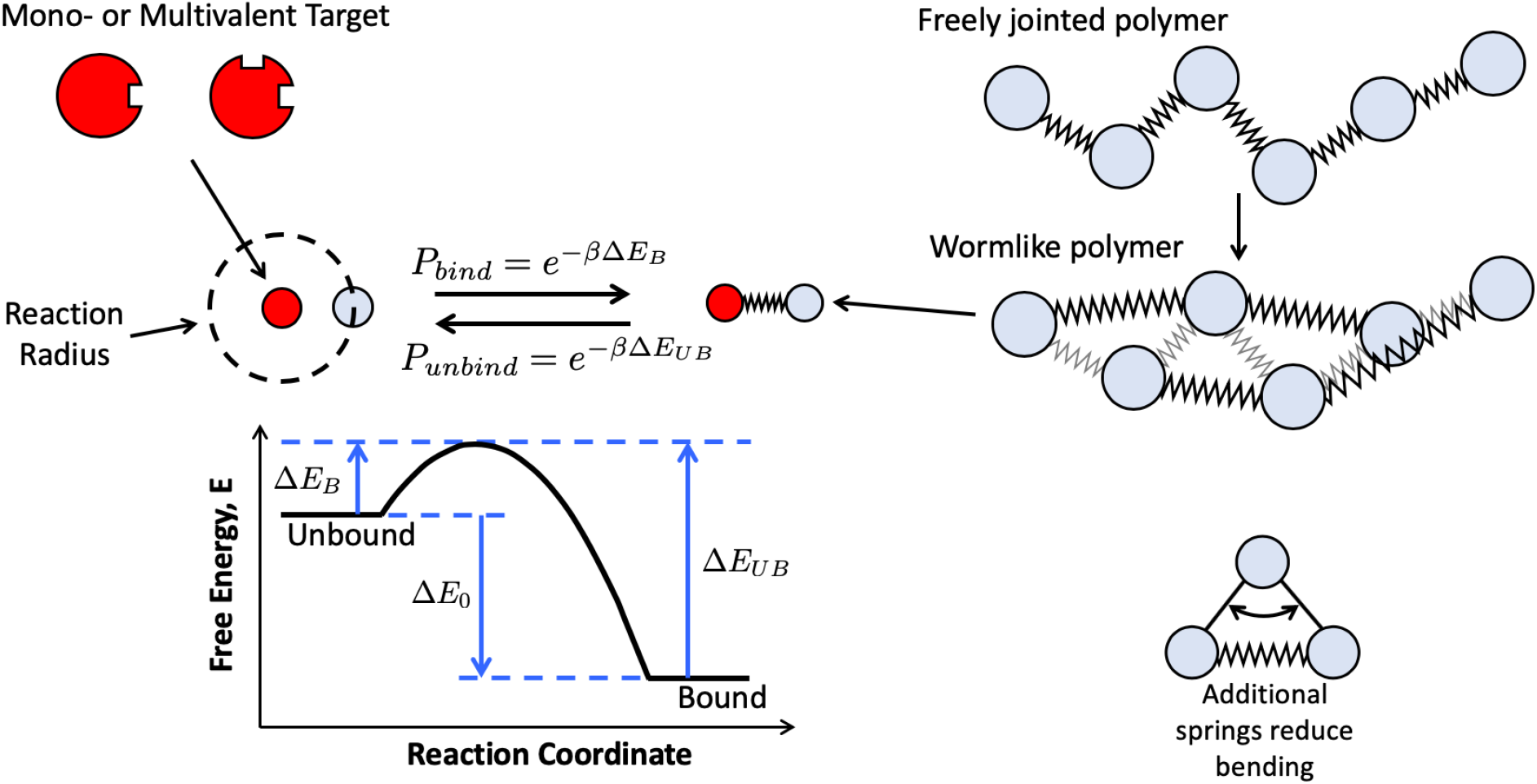
Depiction of simulation scheme. Polymers are represented by spherical beads (light blue) connected by harmonic springs. To introduce stiffness, we employ a simple scheme used also by some commonly utilized force fields (e.g. MARTINI [27]), where an additional spring is placed between every next nearest neighbor along the chain. Each polymer bead has a single ligand, meaning it can only bind monovalently, but making the polymer as a whole multivalent. Targets, on the other hand, can have multiple binding sites and are represented by a single spherical bead (red) with one or two binding sites as shown. Polymer ligands and target binding sites interact when they are within a reaction radius. Within this reaction radius, they have a probability of binding *P*_B_ that depends on the free-energy landscape, as depicted. Once bound, the target and polymer bead are connected by a harmonic spring, and they can unbind with probability, *P*_UB_. Apart from the reactive kinetics that we include here to model the specific binding mechanisms, we use a Lennard-Jones potential to maintain the chain conformation and prevent target-target and target-polymer overlap. This figure is adapted from Zumbro *et al.* with permission from Elsevier [18].

**Figure 2:**
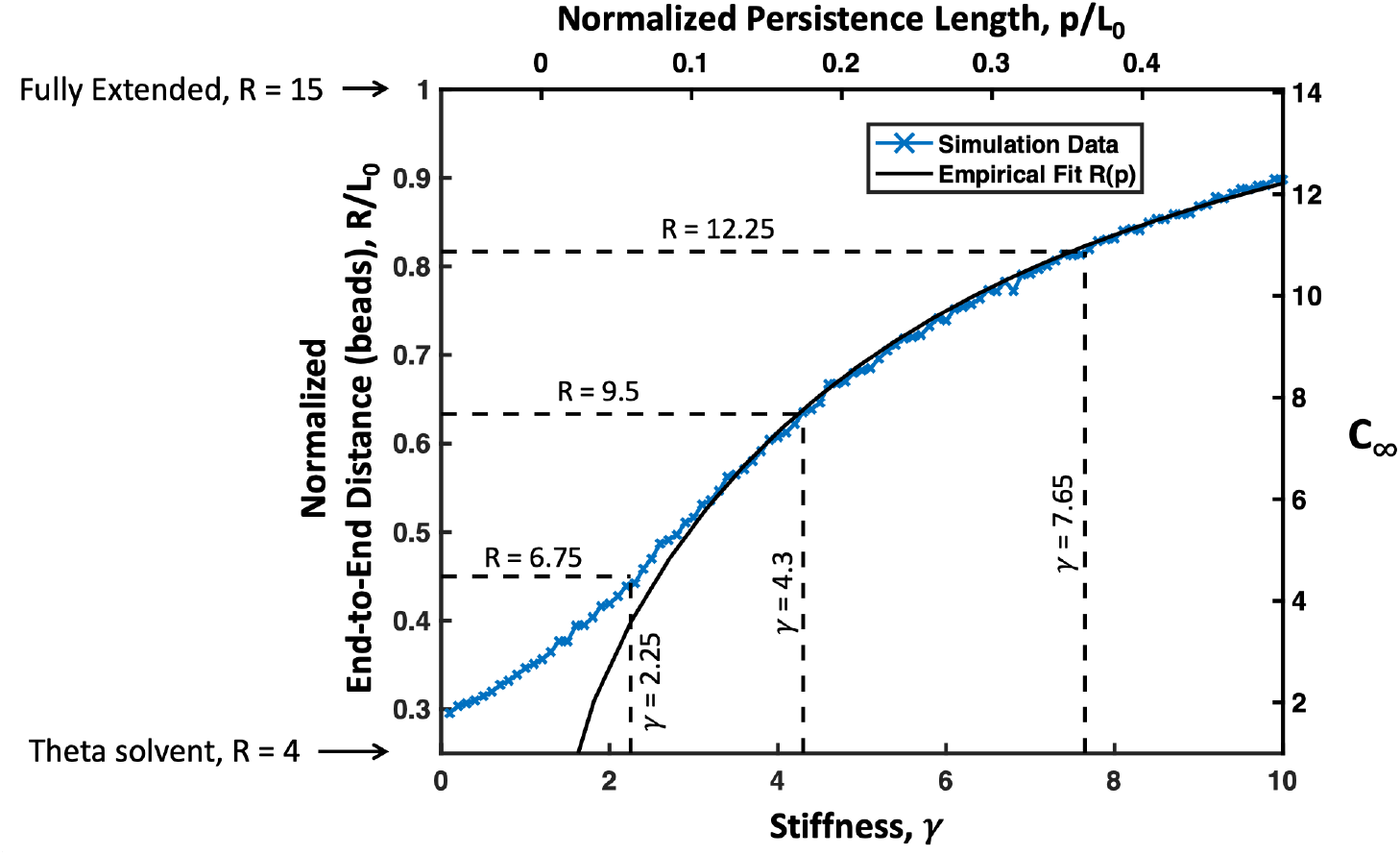
Simulated average end-to-end distance *R* of 16mer polymer chain normalized by the contour length *L*_0_ plotted versus chain stiffness spring coefficient *γ* is shown as blue X’s. Values of *R* and *γ* at which simulations were run are highlighted with dashed lines. These values of *γ* were chosen to explore a wide range of polymer flexibilities and represent the point where *R* ≈ 4 for a perfectly flexible polymer, and 25%, 50%, and 75% of the distance between the most flexible chain *R* ≈ 4 and a perfectly rigid rod where *R* = *L*_0_ = 15. End-to-end distances were converted to *C_∞_* on the right axis and persistence length, *p* on the top axis using the empirical fit relating *R/L*_0_ to *p/L*_0_ (black solid line).

A generic Lennard-Jones potential was applied to control excluded volume and implicit solvation according to:

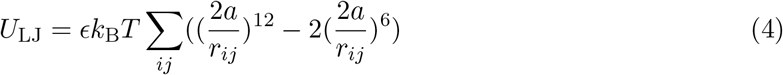

Where *i* and *j* represent two different bead indices and the value of *ϵ* can be adjusted to control the solvent quality and non-specific interactions between beads. Here, we could substitute a screened electrostatic potential but do not expect this to qualitatively change our results. Across the simulations in this work, we have chosen 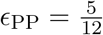 to mimic polymer configurations in a theta solvent [33]. We used polymer target potential 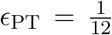 and target-target potential 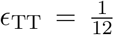 (unless otherwise stated) to mimic a good solvent as summarized in Table 1 Case 1. We chose a theta solvent for the polymer because this is the lower limit in end-to-end distance of a soluble polymer. We would expect a polymer in good solvent to follow similar trends as shown in our previous work, with a smaller range of possible chain end-to-end distances, although this distinction for stiff polymers becomes even more irrelevant.

**Table 1:** *ϵ* values for Polymer-Polymer (PP), Polymer-Target (PT), and Target-Target (TT) bead Lennard-Jones Interactions

| Case # | $\epsilon_{PP}$ | $\epsilon_{PT}$ | $\epsilon_{TT}$ |
| --- | --- | --- | --- |
| 1 | 5/12 | 1/12 | 1/12 |
| 2 | 5/12 | 1/12 | 18/12 |
| 3 | 5/12 | 1/12 | 20/12 |
| 4 | 5/12 | 1/12 | 24/12 |

Our last type of interaction is a reactive lock and key bond, which represents our specific, valence-limited binding interaction. To simulate this reactive binding, harmonic springs were turned on and off between the polymer beads and the targets to dynamically represent bonded and unbonded states. This was implemented using the prefactor Ω(*i, j*) multiplied by a harmonic potential as follows:

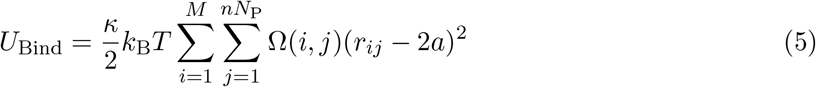

Ω(*i, j*) = 1 when the *i*th binding site on the target is bound to the *j*th bead of the inhibitor, and Ω(*i, j*) = 0 when the target binding site or inhibitor bead is unbound. To control the probability of binding and unbinding, we use a piecewise function based on the energy barriers for the binding reaction from Sing and Alexander-Katz [34].

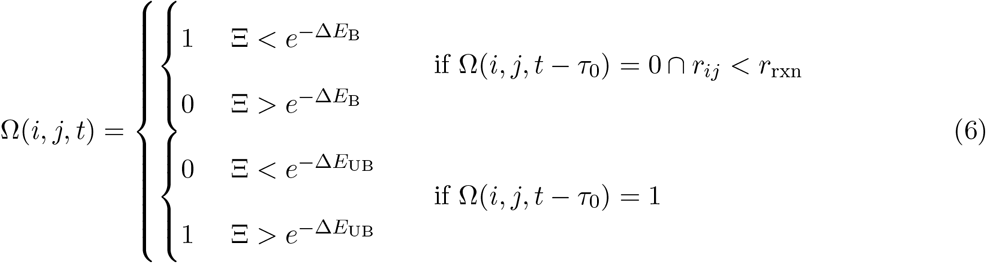

Here, Ξ is a random number between 0 and 1, Δ*E*_B_ is the energy barrier to bind normalized by *k*_B_*T*, and Δ*E*_UB_ is the energy barrier to unbind normalized by *k*_B_*T* as depicted in Figure 1 [34]. Without loss of generality, these energies are considered to be always positive, and the kinetics of binding are held constant by keeping Δ*E*_B_ at 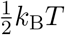 so that binding is a sufficiently frequent event. Increasing or decreasing the energy barrier will respectively slow or accelerate the kinetics of binding and unbinding equally, but not change the system’s thermodynamics. The thermodynamic drive of binding is controlled by varying Δ*E*_0_ = Δ*E*_B_ − Δ*E*_UB_. Binding becomes more favorable as Δ*E*_0_ is made more and more negative. This method is based directly on Sing *et al.* as well as others [34–37]. Binding reactions are evaluated every time interval *τ*_0_ = 100Δ*t*, where Δ*t* is the length of one timestep and *t* is the current time. The reaction radius *r*_rxn_ = 1.1 is the distance apart two beads would be if their surface was touching plus 0.1. Choosing 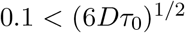 allows time for a target that unbinds to diffuse out of the polymer radius of influence in *τ*_0_ and makes binding events independent [34]. We have applied the constraint that at any time, an inhibitor bead can only bind to one target binding site Σ_*j*_ Ω(*i, j, t*) ≤ 1, and a target site can only be bound to one inhibitor bead Σ_*i*_ Ω(*i, j, t*) ≤ 1. Competing reactions are sampled randomly. Note that we do not include the effect of forces in the breaking of the bonds; this is due to the fact that for forces on the order of *k*_B_*T/a*, this effect is negligible if the characteristic bond length is less than 1 nm. For reference, a discussion of the subject is given by Sing and Alexander-Katz [38].

We used our reactive binding scheme with a free energy of binding per site of Δ*E*_0_ = −4*k*_B_*T*. Each polymer bead contained a single binding site, and each target bead was given one or two binding sites so that we could compare the monovalent case to the divalent case. Assuming the size of a target bead to be approximately 5 nm in diameter, and using langmuir adsorption theory, we can convert this Δ*E*_0_ = −4*k*_B_*T* binding energy into a dissociation constant in Molar, resulting in a monovalent binding affinity of *K*_D_ = 0.1 mM. Details of this conversion are shown in the supplemental information. This monovalent binding affinity is well within the weakly binding mM to *μ*M range typical of lectins and sugars binding as well as the mM to *μ*M affinity range of some monovalent protein-protein and RNA-protein binding found in biocondensates [19, 39–42].

The potentials are applied over the timestep 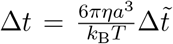 where 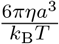 is the characteristic monomer diffusion time or the time that it takes a bead to diffuse its radius *a* and the dimensionless timestep is 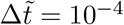. These equations are all made dimensionless by scaling energies by thermal energy *k*_B_*T*, lengths by bead radius *a*, and times by the characteristic diffusion time 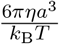.

The simulation code is available by request.

## Results and Discussion

For all simulations we placed four polymers with a degree of polymerization *N*_P_ = 16 because this is slightly above the length at which increasing polymer length leads to only a small increase in binding affinity for a perfectly flexible polymer chain binding to a divalent target [18]. The binding dependence on length for flexible polymers is primarily influenced by the loop sizes that can favorably occur when bonded twice to a target. Thus, we expect that in wormlike chains, intra-chain loops will be even shorter, and so overall polymer length will be relatively unimportant. Simulations for both the dilute case of a single target binding to our polymers as well as the high concentration case where many targets interact with the polymer simultaneously are discussed in this work.

### Binding to a single target

It is important to consider how a single mono or multivalent target binds to a polymer without competition from other targets for available binding sites. We did this by placing a single target with either one or two binding sites in with four identical 16mer polymers as shown in Figure 3A. To examine the binding affinity of our polymers we calculated the average time bound *τ*_B_ for our target, where we considered our target bound whenever at least one of its binding sites was bound to the polymer. Accordingly, we consider the target unbound whenever none of its binding sites were bound to the polymer and denote the average time interval unbound as *τ*_UB_.

**Figure 3:**
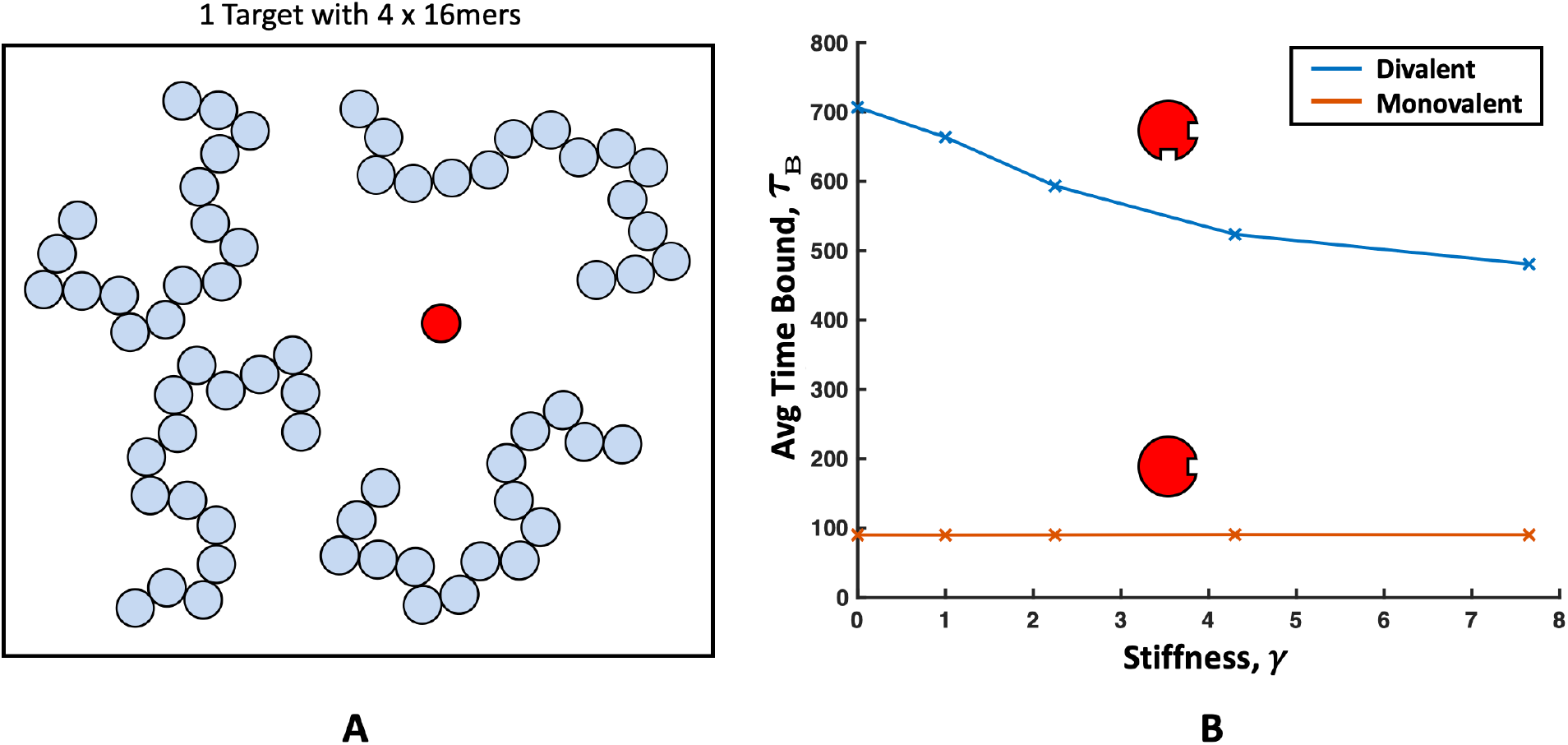
(A) Schematic of the single target simulation set up with a single mono- or divalent target shown in red and four polymers with a length of 16 beads. (B) The average time interval bound *τ*_B_ of a single divalent target (blue) and a monovalent target (orange) versus the polymer stiffness controlled by the angle-bending spring coefficient *γ*. Higher *γ* corresponds to stiffer springs and more rigid polymers. The monovalent target *τ*_B_ seems unaffected by the polymer chain stiffness while the divalent targets show a decrease in *τ*_B_ with *γ*.

We have plotted the *τ*_B_ for for divalent and monovalent targets in Figure 3B. From this plot, we can see that the *τ*_B_ for the monovalent target does not depend on polymer stiffness, but the *τ*_B_ of a divalent target decreases with increasing bending spring coefficient *γ*. Remember that higher *γ* corresponds to a stiffer polymer. While this decrease was predicted by Zumbro *et al.*, the previous theory for random coil chains only partially applies [18]. Previous work showed that the entropic cost of forming loops limits the polymer binding affinity, but in the case of a rod-like polymer or worm-like chain, loop entropy is not the limiting factor. Instead, we predict that the major energetic factor limiting multivalent binding affinity is the enthalpic cost of bending the polymer to make two contacts with the target. If bending is the limiting factor, we can estimate the energy as:

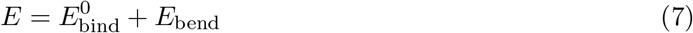

where 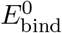 is a constant denoting the favorable energy of forming two bonds while *E*_bend_ is the quantity that is changing most drastically with loop length. When the target is bound twice to the same polymer chain, once to polymer bead *i* and simultaneously once to polymer bead *j*, we define the loop length by subtracting the two values *l*_loop_ = |*i* − *j*|. This results in *l*_loop_ = 1 when *i* and *j* are right next to each other, *l*_loop_ = 2 when there is a single unbound bead between them, etc. To minimize bending penalty for a particular *l*_loop_, the polymer will want to minimize curvature or maximize the bending radius *R*_loop_. If we consider very small loops where *l*_loop_ = 1, 2, 3, the maximum *R*_loop_ is constant at 2*a* where *a* is the radius of the beads. Because our target is very coarse-grained, we allow its binding sites to be accessible anywhere on the surface; this makes the largest *R*_loop_ for *l*_loop_ = 1, 2, 3 occur when the polymer beads all exactly touch the surface of the target. Since at small *l*_loop_, *R*_loop_ is constant, we can estimate the probability of forming a loop as:

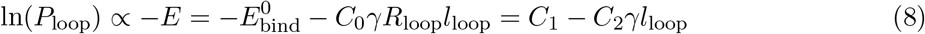

where *C*_0_ is a constant representing the bending cross-section, and where, for small loops, we have replaced 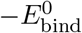 and *C*_0_*R*_loop_ with generic constants *C*_1_ and *C*_2_ respectively. From Equation 8, we find that if bending energy is the most significant influence on *τ*_B_, we should see the ln of the frequency of increasing loop size decay linearly with a rate proportional to the chain stiffness *γ*.

We can probe this theory directly by examining the length of loops formed and their frequency depicted in Figure 4. From this plot, we can see that the more flexible polymers (*γ* = 0, 1, or 2.25) follow an exponential decay characteristic of entropic loop costs [18], while the stiffer chains (*γ* = or 7.65) follow a linear decay in loop length characteristic of an enthalpic bending loop cost. This cost increases as *C_∞_* increases, leading to a drastic drop off in loop lengths leading to only *l*_loop_ ≤ 2 being formed more than 1% of the time for *γ* = 4.3 and 7.65. Since only small loops are formed for highly stiff polymers, we can test our theory on them by measuring their slope.

**Figure 4:**
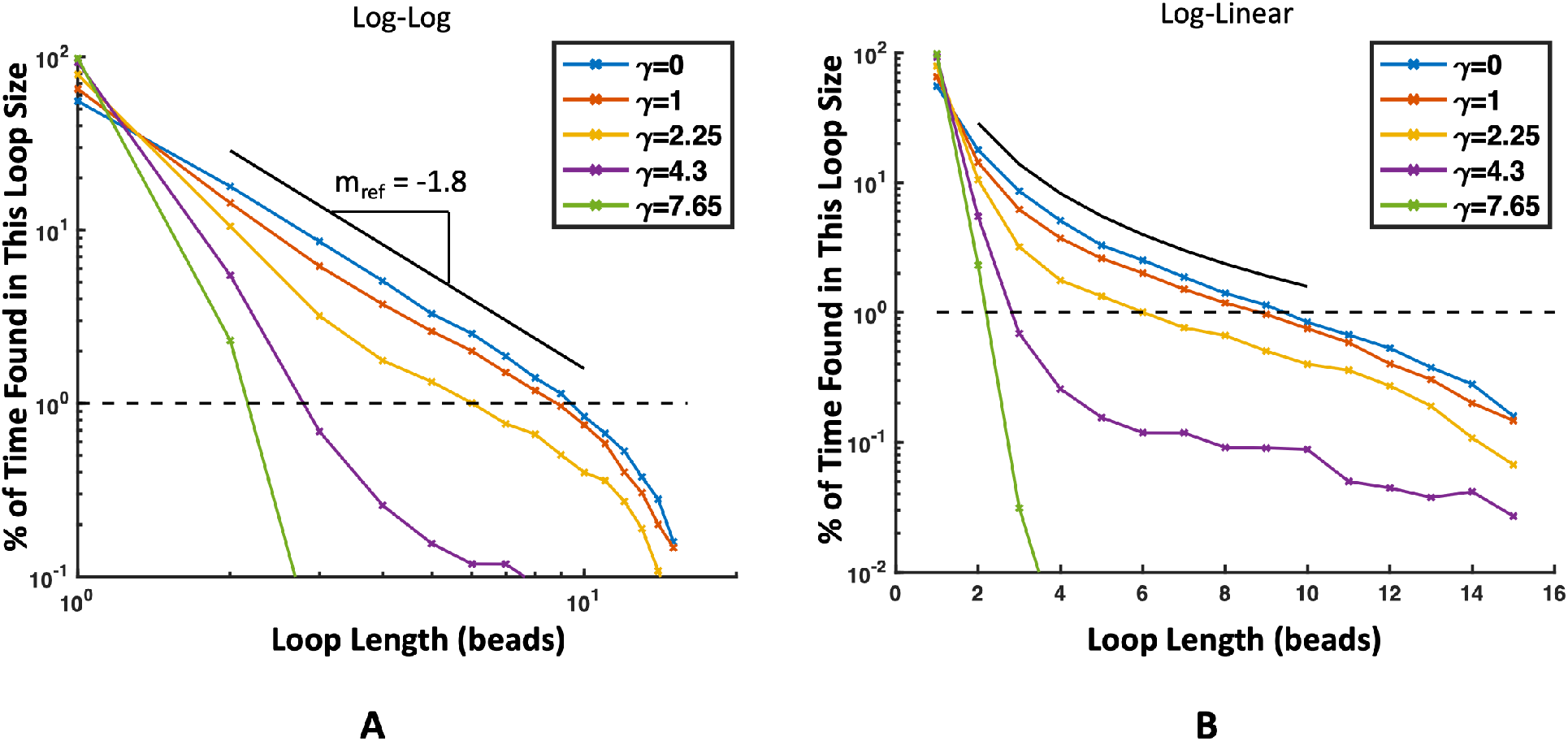
Percent of time that a polymer bound twice to a target is in a certain loop length plotted in (A) log-log scale and (B) log-linear scale. Each color represents a different polymer stiffness, the dashed black line represents 1%, and the solid black line is an example of *y* = *x^−^*^1.8^. The frequency of long loops decreases as polymer stiffness increases. (A) More flexible chains (*γ* = 0, 1) have a power law decay in loop size due to the entropic cost of forming loops [18]. This manifests as a straight line in the log-log scale. (B) Stiffer chains (*γ* = 4.3, 7.65) have an exponential decay in loop lengths for short loops due to the energetic cost of bending. We can see this manifest in the log-linear plot as a straight line for short loop lengths (*l*_loop_ = 1, 2, 3). Lines are for aiding the eye and are not a theoretical fit.

Fitted values for *C*_1_ and *C*_2_ from Equation 8 are shown in Table 2 with corresponding bending theory lines shown with simulation data in Figure 5. For the values shown in Table 2, we have chosen to show the Y-intercept at *l*_loop_ = 1 because this corresponds to the case of 0 angular springs between bound polymer beads *i* and *j* and because a loop length of 0 is nonsensical in this context. From these best fit lines, we find an excellent fit for such a simple theory for the stiffest polymer (*γ* = 7.65) with worsening fit as *γ* decreases, likely due to increasing contributions from entropic loop costs dominating flexible polymers. As predicted, the value of *C*_1_, which represents the binding energy unaffected by chain stiffness is almost constant for the three highest stiffness chains. The small differences in *C*_1_ are likely due to some dependence of 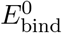 on *γ* which our model does not capture such as increased stress on target-polymer bonds in stiffer chains, and unbound polymer beads in the center of the loop pressing in toward the target and into its excluded volume. Both of these secondary effects are energetically unfavorable and would lead to an increase in 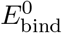 with *γ*, which aligns well with our calculated values of *C*_1_.

**Table 2:** Slopes and intercepts for lines fitted to loop lengths for Eq. 8 and plotted in Figure 5.

| $\gamma$ | Y-intercept ($C_1$) | Slope Constant ($C_2$) |
| --- | --- | --- |
| 2.25 | 4.23 | -0.71 |
| 4.30 | 4.41 | -0.57 |
| 7.65 | 4.66 | -0.52 |

**Figure 5:**
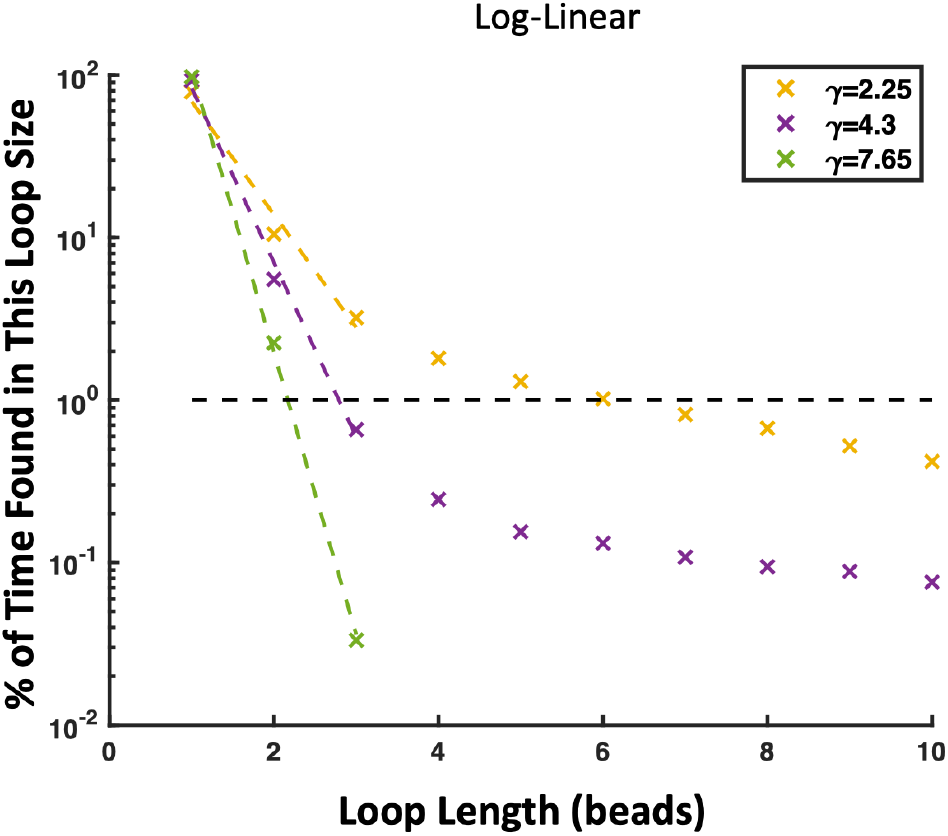
Frequency of loop sizes in Log-Linear scaling. ‘x’s denote simulation data and dashed (- - -) lines represent the best linear fit following equation 8 with values of *C*_1_ and *C*_2_ listed in Table 2.

As for the slope coefficient *C*_2_, the behavior of the chain with *γ* = 2.25 is the least well-captured. This is likely because the chain is still relatively flexible and experiencing a blend of the bending costs of wormlike chains and the entropic cost of freely jointed chains. In contrast, we found good agreement for the two stiffest chains (*γ* = 4.3, 7.65) with our estimate that the decay rate of loop length should be constant *C*_2_ times the chain stiffness *γ*. The fit *C*_2_ values for the two stiffest chains are within 10% of each other. Thus, with this very simple theory of bending, we capture quite well the behavior of stiff chains.

As a result, we conclude that in the case of a much larger polymer binding to a single multivalent target, stiff polymers are limited by the enthalpic cost of bending when forming an intra-polymer loop. Because long loops cost high amounts of bending energy, they are almost impossible to form, resulting in fewer possible binding arrangements for the target and overall decreasing the *τ*_B_ of the rigid polymer over a flexible polymer. We can extend our results from *τ*_B_ to relative overall binding affinity by also measuring the average time unbound *τ*_UB_ and calculating the relative dissociation constant as 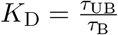 with measured *τ*_UB_ and *K*_D_ shown in Figure 6A and B respectively.

**Figure 6:**
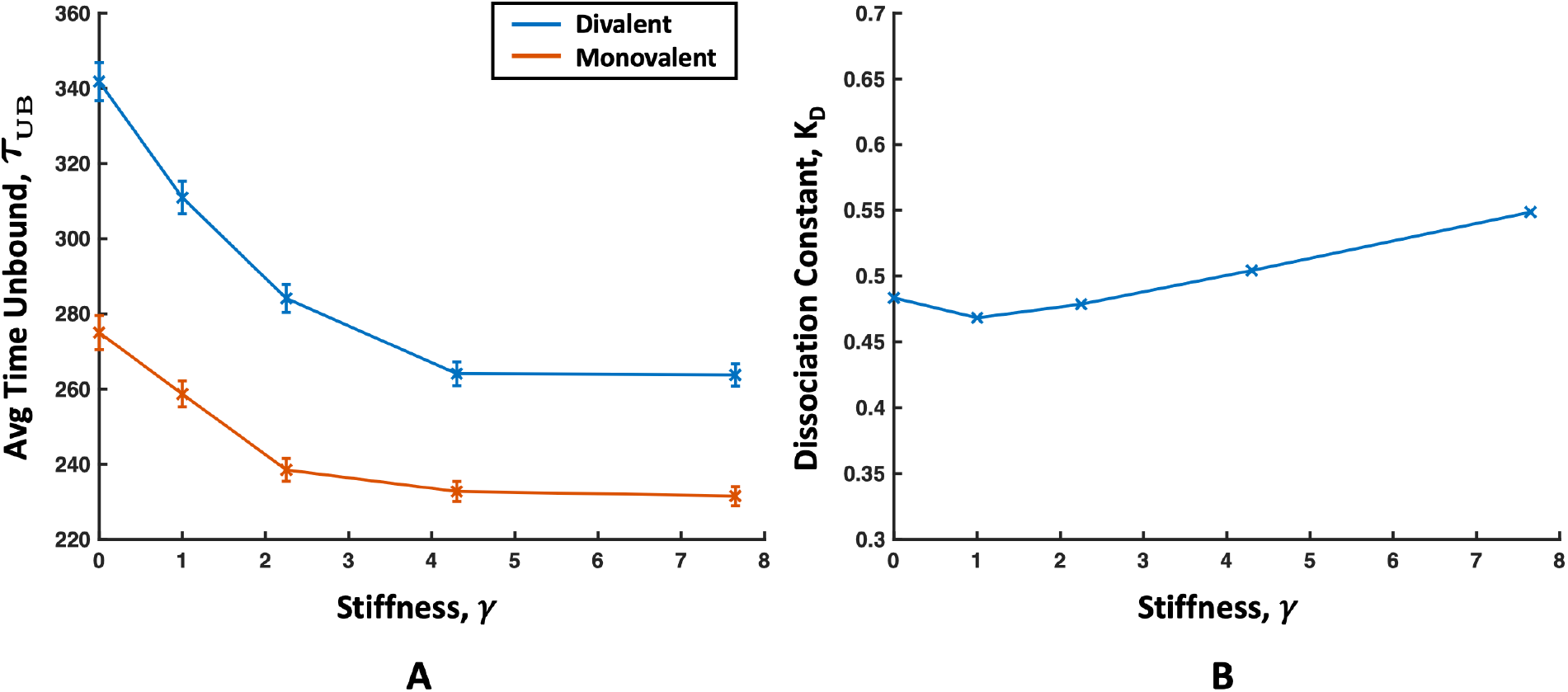
(A) The average time interval unbound *τ*_UB_ for a single mono- (orange) or divalent (blue) target binding to a polymer. The *τ*_UB_ decreases similarly for both target valencies because it is dependent on the distribution of polymer binding sites throughout the simulation volume. Standard error is denoted by error bars. (B) Dissociation constant *K*_D_ for a divalent target versus polymer stiffness. The longer *τ*_UB_ is not enough to overcome the longer *τ*_B_ for flexible polymers and flexible polymers show a lower *K*_D_ (higher affinity) than rigid ones. A plot of the *K*_D_ for a monovalent target is dominated by *τ*_UB_ and is shown in Figure S4.

The *τ*_UB_ is a combination of the time it takes for a target to come within reach of a free polymer binding site, which is dependent on the morphology of the polymer, multiplied with the binding attempt rate and success rate, which are constant for our simulations. By approximating the flexible polymer as a sphere and the rigid polymer as a thin cylinder, we found that the diffusive flux of targets toward the sphere is smaller than a cylinder for our polymer concentration. We can think of this as the more rigid polymers having binding sites more uniformly distributed throughout the volume. This means that on average, targets take longer to find flexible polymers than rigid ones and the *τ*_UB_ shortens with *γ*. Diluting the polymer concentration can flip the relationship between the target flux toward a cylinder and sphere so that 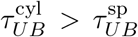, but this will only exaggerate the effect of stiffness on *K*_D_ plotted in Figure 6B. A detailed discussion of the flux calculation are presented in the supplemental information in Section 2, Figure S2.

While *τ*_UB_ decreases with increasing chain stiffness shown in Figure 6A, the decrease is not enough to overcome the decrease in *τ*_B_ with chain stiffness. This results in a smaller dissociation constant *K*_D_ for stiffer chains. The effect is not monotonic, with a slight decrease in the *K*_D_ as *γ* increases from 0 to 1, which is probably due to increased free volume and therefore more accessible binding sites as seen in an experiment with polymers binding to lectins [6]. But as we move away from slightly extended chains that still follow a random walk to highly extended chains with end-to-end distance much greater than, there is a distinct upward trend in 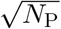. Since smaller *K*_D_ corresponds to a stickier polymer, we find that the affinity of the polymer generally decreases with increasing *C_∞_*, and conclude that this result is primarily affected by the transition of the binding regime from a flexible chain, where the entropic cost of loop formation dominates, to the regime of a wormlike chain where the high enthalpic cost of bending to create divalent loops dominates and reduces binding affinity.

Our results do not contradict previous research on precise ligand engineering where it was found that for matched sizes on the size scale of the target, stiff linkers have higher affinity [2, 5, 8, 13, 14]. Conversely, our research is complementary to previous works; with our result of large flexible polymers showing higher binding affinity than large rigid polymers, this suggests that the most sticky polymer might consist of flexible regions between the stiff, matched size binding sites detailed in the aforementioned studies.

### Binding to multiple targets

In this section, we consider the binding of many targets to polymers with varying backbone flexibility as shown in Figure 7. In this case, we place 64 mono or divalent targets in with our four 16mer polymers to create competition and examine the conditions under which a condensed phase might nucleate with the help of the polymer. 64 targets corresponds to the concentration at which the number of target beads matches the number of polymer beads and there is significant competition. For monovalent targets, the number of binding sites on the targets is the same as on the polymer while the total number of binding sites for 64 divalent targets is twice the number of polymer binding sites. We can consider the concentration of targets in real units by assuming a target diameter. For example, assuming a target protein diameter of 5 nm, 64 targets corresponds to 100 *μ*M and assuming a weight of 70 kDa, approximately 7 mg/ml.

**Figure 7:**
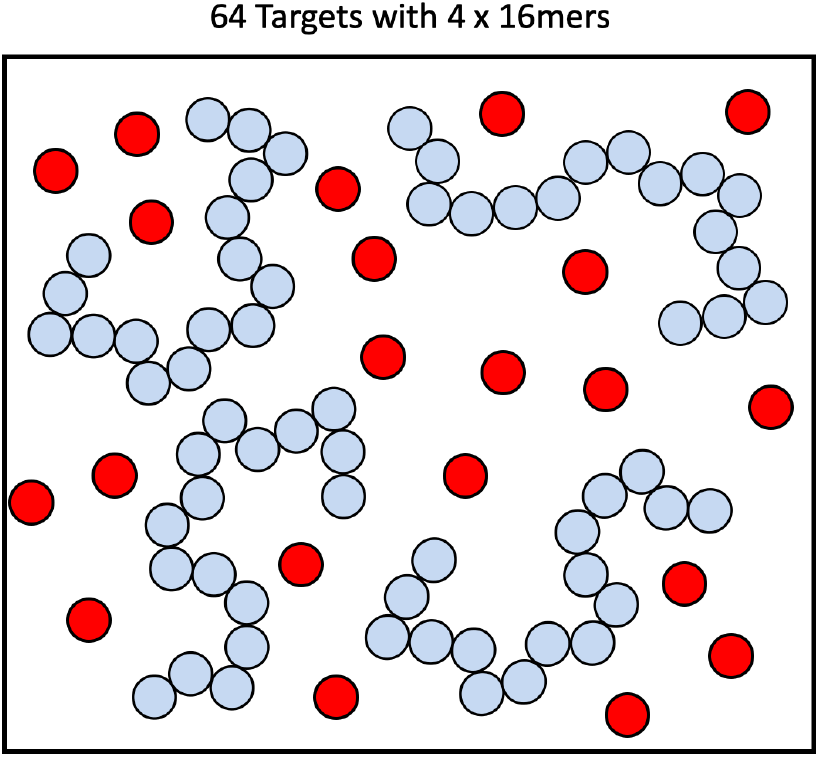
Schematic of simulations with multiple targets. In this case, 64 targets are placed in a box with 4 16mer polymers to examine how target-target interactions and competition between targets for binding sites on the polymer can change the phase behavior of the system.

Because we have multiple targets, we now need to consider the interactions between targets. To capture these non-specific interactions that control protein solubility limits we add a generic Lennard-Jones potential between the targets themselves as described in Eq. 4. We used several values of potential well energies ranging from almost no attraction (*ϵ*_TT_ = 1/12) to moderate levels of attraction (*ϵ*_TT_ = 24/12) to explore the wide range of solubilities found in proteins. Parameters for the cases studied are summarized in Table 1. By themselves, the 64 targets are soluble throughout this entire range of intra-target interaction strengths, and only condense on their own at approximately *ϵ*_TT_ = 30/12 or 2.5*k*_B_*T* Lennard-Jones attraction shown in a rough phase diagram in Figure 9A.

Previous work has shown that increasing polymer length can induce a condensed phase, so we wanted to further investigate how changing the polymer backbone stiffness changes the phase boundary [18]. In doing so, we hope to provide insights for research on liquid-liquid phase separation as well as those targeting inhibition of high concentrations of multivalent toxins. To this end, we used visual inspection to look for a persistent condensed phase in our simulations. We considered our systems “phase separated” if the targets and polymers formed a condensed globule that persisted throughout the second half of the simulation time and “not phase separated” or “mixed” if no globule formed or if the globule was unstable and repeatedly broke apart during the simulation time. The results of this inspection were compiled into a phase diagram in Figure 8B with renderings of specific cases provided for reference.

**Figure 8:**
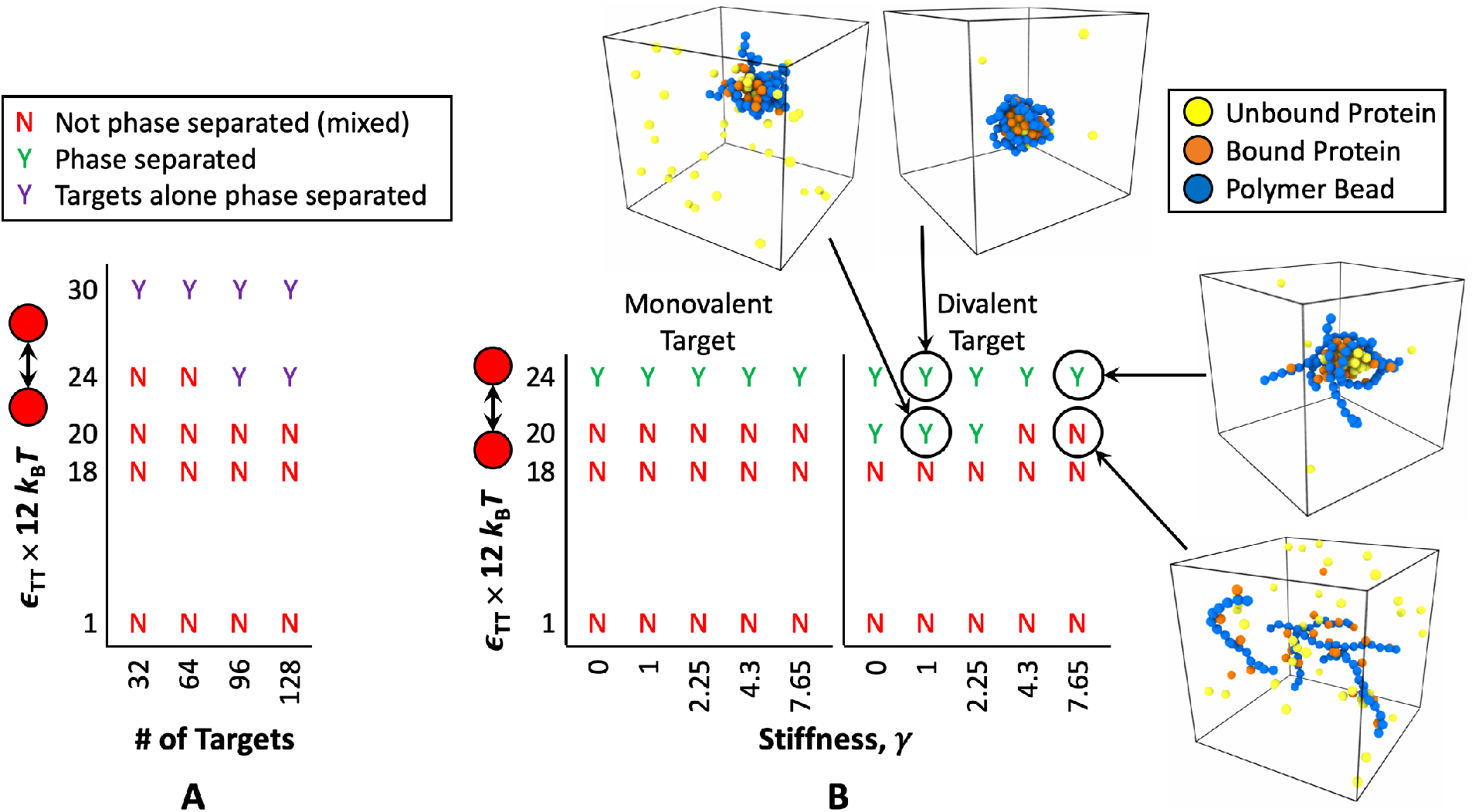
Phase diagrams of (A) targets only with increasing concentration of targets on one axis and increasing target-target Lennard-Jones attraction on the other and (B) 64 target and 4 16mer polymer simulations with polymer stiffness on one axis and target-target Lennard-Jones attraction on the other. Results are shown for both mono and divalent targets. A not phase separated or “mixed” system is denoted by a red letter “N” for “no”, a phase separated system where the polymer and targets are both components of the condensed phase is denoted by a green “Y” for “yes”, and a purple “Y” denotes a system where the targets phase separated by themselves, in this case because no polymer was added. (A) Shows that at a 64 target concentration, the targets by themselves (without the addition of polymer) are soluble at all Lennard-Jones attractions studied in the polymer-target systems in (B).

From these phase diagrams we find that, first, just the addition of the polymers lowers the phase boundary for 64 targets below *ϵ*_TT_ = 24/12 for all polymer flexibilities and target valencies. Additionally, increasing the valency of the target lowers the phase boundary, consistent with previous research on liquid-liquid phase separation of multivalent polymers [19, 20, 23]. Looking at the renderings of simulations with divalent targets, it is clear that even though both systems are phase separated at *ϵ*_TT_ = 24/12 attraction, the resulting complexes look very different depending on the chain stiffness. For flexible polymers *γ* = 1, the resulting complex is spherical, characteristic of a liquid globule, and the polymers are coating the surface relatively tightly. In the case of the stiffest polymer *γ* = 7.65, we still see a rounded globule of targets, but now the polymer is not completely stuck to the condensed target surface. Instead, stiff polymers have peeled off the globule and are sticking out making the complex look spiky or hairy. Because these stiff chain ends are sticking away from the condensed target phase, we expect that they are less bound, making the *K*_D_ larger (lower affinity) for the targets and lowering the number of total sites occupied on the polymer. These effects lower the binding efficiency of the stiff polymers and, in the case of multivalent targets, result in the phase boundary being pushed to higher target-target attractions as the polymer stiffens. This appears as the solubility limit of our divalent targets being pushed to higher energies from below *ϵ*_TT_ = 20/12 to *ϵ*_TT_ = 24/12 in Figure 8B. This phenomenon has also been seen in complex coacervates where it was shown that the two phase region shrinks as the stiffness of the binding polycation/polyanion species increases [43].

In Figure 9A, we have plotted the dissociation constant for divalent targets as 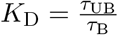 vs *γ* for several values of *ϵ*_TT_ with the values of *τ*_B_, *τ*_UB_, and monovalent *K*_D_s plotted in the supplemental information Figure S4. As expected from the earlier single target case and our phase diagram, intratarget attraction and chain stiffness have a significant effect on the binding of divalent targets. As target-target attraction increases, the *K*_D_ decreases for all polymer stiffnesses. This is because when many targets are bound to the polymer, bound targets benefit energetically from being near other neighbors also bound to the polymer chain. This additional energy benefit makes the polymer appear stickier and is magnified when the polymer can nucleate a condensed target phase because then bound targets can gain the energy benefit of being near both bound neighbors and unbound neighbors. This can be seen in Figure 9A by looking at the values of *K*_D_ for intra-target attraction *ϵ*_TT_ = 20/12. Flexible polymers *γ* = 0, 1, 2.25 have a low and almost constant *K*_D_ because the polymers nucleated a condensed target phase and so their binding affinity is benefiting greatly from the favorable energy of clustered unbound targets. As *γ* increases from 2.25 to 4.3 we see a sharp increase in the *K*_D_, due to the fact that phase separation is harder to induce with a stiff polymer. Subsequently, targets that bind to a rigid polymer will have fewer target neighbors and benefit less from favorable target-target energies.

**Figure 9:**
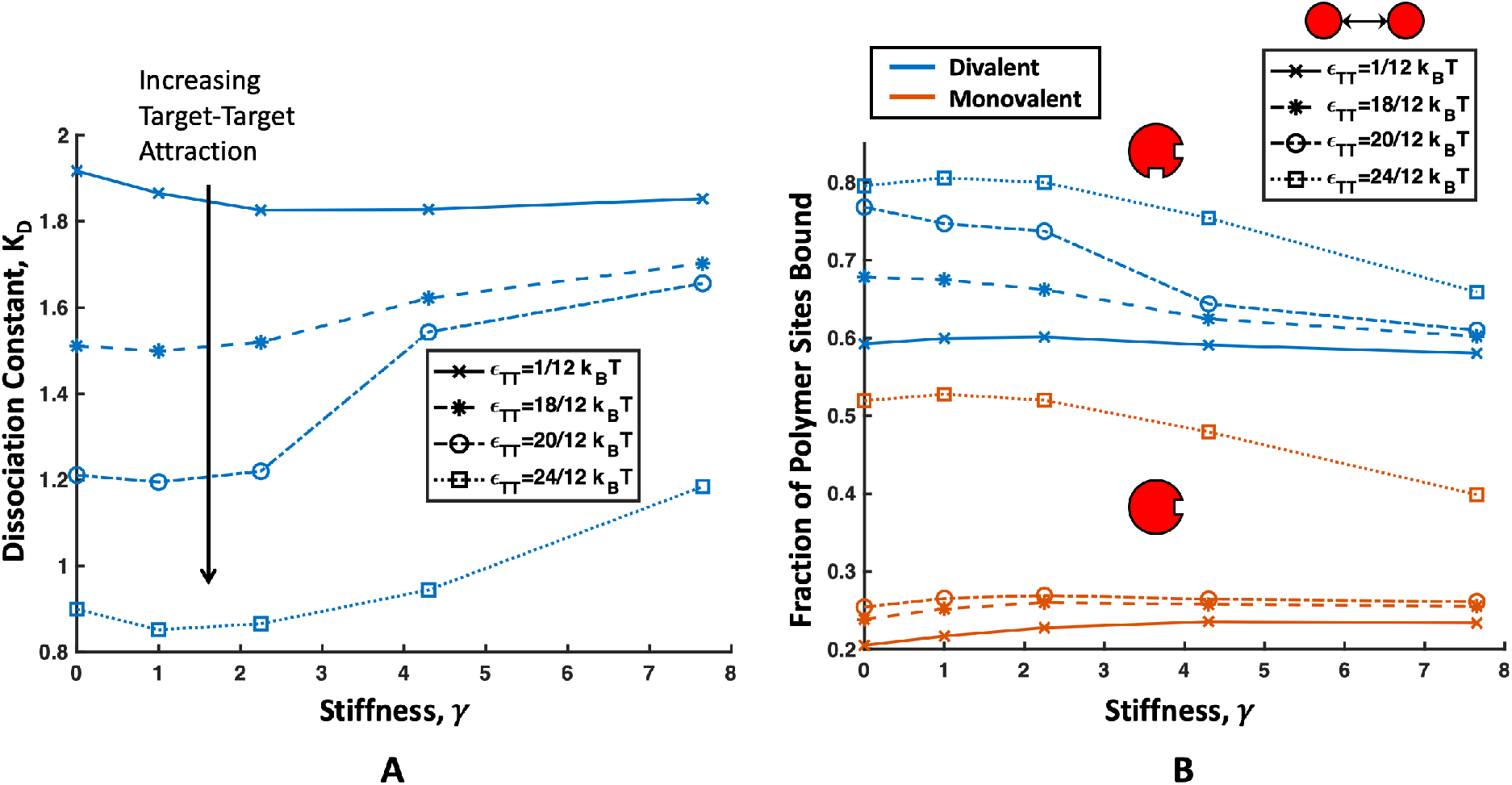
(A) *K*_D_ for 64 divalent targets binding to 4 16mer polymers. As target-target attraction *ϵ*_TT_ increases, *K*_D_ decreases. For *ϵ*_TT_ = 1, binding affinity is dominated by the increased *τ*_UB_ and flexible polymers are slightly higher affinity than stiff ones. At *ϵ*_TT_ ≥ 18/12, *τ*_B_ dominates and flexible polymers have higher affinity than stiff polymers. We can see a sharp increase in *K*_D_ for *ϵ*_TT_ = 20/12 as *γ* increases from 2.25 to 4.3 signaling the phase boundary where flexible polymers are able to nucleate a condensed target phase but stiff polymers are not. (B) Binding efficiency of polymers calculated as the average fraction of sites on the polymer bound versus *γ*. This plot closely mimics the one for *K*_D_, with a sharp decrease in binding efficiency for divalent targets at *ϵ*_TT_ = 20/12 denoting the phase transition. For phase separated systems at *ϵ*_TT_ = 24/12, there is an approximately 10% decrease in sites bound on the polymer between the *γ* = 0 and *γ* = 7.65 for both target valencies. This is due to rigid polymer resistance to bending and their tails sticking out away from the condensed phase as shown in Figure 8B.

We can confirm this reasoning by directly examining the clustering of unbound targets near the polymers. We plot the radial distribution function (RDF) of unbound targets for all simulation conditions in Figure 10. While general curve shapes for *ϵ*_TT_ = 1/12, 18/12, and 24/12 show only small changes due to polymer stiffness, the profile of unbound targets near the phase boundary *ϵ*_TT_ = 20/12 is strongly dependent on polymer stiffness. From the RDFs of unbound targets at *ϵ*_TT_ = 20/12 (Figure 10B), we can see that the flexible polymers are able to stabilize unbound targets at this target solubility while stiff polymers are not. In order for stiff polymers to be able to form a stable cluster of unbound targets, the target-target attraction must be increased to *ϵ*_TT_ = 24/12. Even when phase separation occurs at *ϵ*_TT_ = 24/12 in Figure 10C, the stiffer polymers stabilize fewer unbound targets shown by the lower RDF, making them lower affinity than their flexible counterparts.

**Figure 10:**
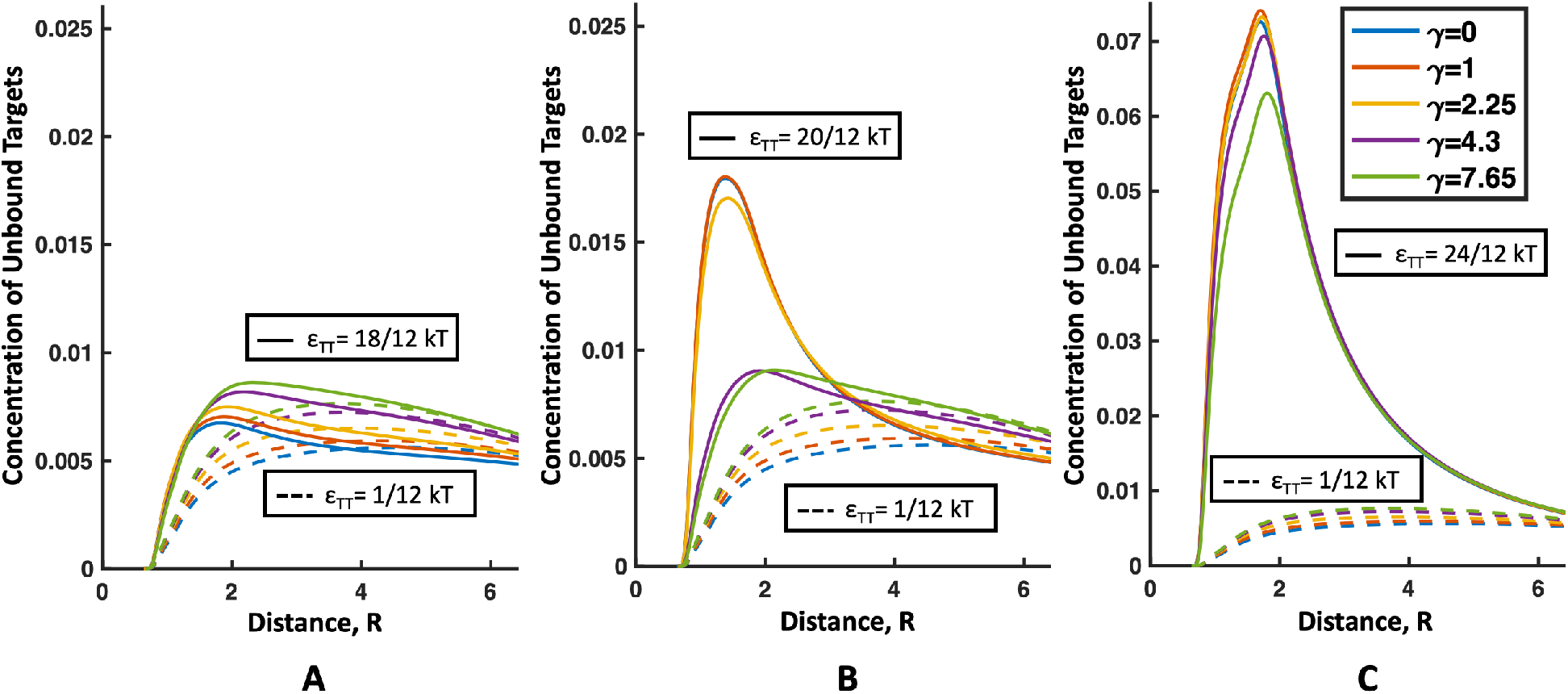
Radial distribution function (RDF) for concentration of unbound targets found near the polymer chain where the x-axis R is the distance from a target center of the closest polymer bead. In all plots, the dashed line represents almost no target-target interaction *ϵ*_TT_ = 1/12. (A) The solid lines represent the RDF of targets for *ϵ*_TT_ = 18/12. Only small changes in the RDF occur with stiffness. (B) The solid lines represent the RDF of targets for *ϵ*_TT_ = 20/12. Note that the solid red (*γ* = 1) and solid blue (*γ* = 0) lines overlap. Here, flexible polymers show a much higher concentration of unbound targets near the chain because they are able to induce phase separation at this target-target potential. (C) The solid lines represent the RDF of targets for *ϵ*_TT_ = 24/12. Note that the blue, red, and yellow lines overlap (*γ* = 0, 1, 2.25). All polymers cause phase separation at this *ϵ*_TT_, so all flexibilities show increased concentration of unbound targets near the polymer.

We can also look at the polymer binding efficiency shown in Figure 9B to understand why stiff polymers phase separate at higher target-target attractions than flexible polymers. Here we can see that for *ϵ*_TT_ = 24/12, where all polymers nucleate phase separation, the percent of polymer sites occupied drops more than 10% as *γ* goes from 0 to 7.65. This is likely due to the hairy or spiky ends shown in the rendering of stiff chains in Figure 8B. These stiff chain ends that extend away from the condensed phase rarely get to interact with the target globule since they would have to bend in to access them, and bending for stiff chains is energetically costly. This results in the parts of the chain extending away from the globule being relatively unbound and decreasing the binding efficiency of the whole polymer. Since the rigid ends are extended out into the dilute target phase, they interact with targets more like our dilute target case discussed above, where they are already lower affinity than flexible chains due to the costs of bending associated with polymer loops. Therefore, resistance to bending makes stiff polymers bind less efficiently, making it harder to collect the critical concentration of bound targets needed to stabilize and collect unbound targets. This explains the change in the phase boundary to higher *ϵ*_TT_ as *C_∞_* increases.

Overall we have found that the phase boundary between a gas-like and condensed target/polymer phase depends on the polymer’s *C_∞_*. Flexible random-coil polymers can lower the solubilities of target proteins more significantly than rigid wormlike polymers, and thus we expect that modulating polymer stiffness could play a role in controlling the phase separation of synthetic systems as well as biological liquid-liquid phase separation.

## Conclusion

In this work, we have presented on how the flexibility of multivalent polymers influences binding to much smaller targets. When binding to dilute targets where there is little competition for binding sites, we have shown that there are two multivalent binding regimes. Flexible random coil polymers fall into the first regime, where binding twice to the target is dominated by the loss of entropy of the the polymer loop as described earlier by Zumbro *et al.* [18]. We have shown here that stiff polymers fall into a second regime where binding affinity is dominated by the enthalpic cost of bending into a loop when binding divalently to a target. The high cost of bending makes stiff rod-like polymers have lower binding affinity for targets than random coil polymers. Therefore, combined with previous research showing that rigid molecules with precisely spaced binding sites have the highest affinity for targets of similar size, we expect that long polymers should ideally contain small rigid binding sections with flexible linkers connecting them into a larger chain in order to achieve the highest affinity [2, 5, 8, 13, 14].

Next, we extended our simulations to the case of many targets binding to the polymer at the same time. This adds competition between our targets as well as allowing us to consider the non-specific interactions between the targets themselves. We show that the presence of polymers can lower the solubility limit of the targets for both multivalent and monovalent binding. In the case of divalent binding we show that flexible polymers can nucleate a condensed phase at lower intra-target attractions than rigid polymers. The morphology of the resultant condensate is different for flexible and rigid polymers with flexible polymers creating a relatively smooth ball and with rigid polymer tails sticking out away from the spherical droplet of condensed targets. This resistance to bending, lowers the binding efficiency of the rigid polymers and makes it more difficult for them to nucleate a condensed phase. Since more flexible polymers phase separate at lower energies, perhaps the drastic changes possible in DNA flexibility could have implications on DNA-containing biocondensate formation. While more investigation is needed on this topic, a recent experimental study showed that more flexible DNA favors liquid-liquid phase separation [44].

We hope our results will aid in the design of new polymeric toxin inhibitors as well as help scientists better understand the formation of membraneless organelles and how changes in polymer stiffness can modify the phase boundary of biocondensates.

## Author Contributions

- Designed research (EZ, AAK)
- Performed research (EZ, AAK)
- Analyzed data (EZ, AAK)
- Wrote the manuscript (EZ, AAK)

## Acknowledgments

The authors were supported by the Department of Defense (DoD) through the National Defense Science and Engineering Graduate (NDSEG) Fellowship Program. The authors were also supported by the Ida M. Green Fellowship and the Collamore-Rogers Fellowship through the MIT Office of the Dean of Graduate Education.

## Supporting Information

The following files are available free of charge.

SupportingInformation.pdf: Additional calculations and plots Figures S1-S5.

